# Microsaccades during high speed continuous visual search

**DOI:** 10.1101/2020.04.28.062042

**Authors:** Jacob G. Martin, Charles E. Davis, Maximilian Riesenhuber, Simon J. Thorpe

## Abstract

Here, we provide an analysis of the microsaccades that occurred during continuous visual search and targeting of small faces that we pasted either into cluttered background photos or into a simple gray background. Subjects continuously used their eyes to target singular 3-degree upright or inverted faces in changing scenes. As soon as the participant’s gaze reached the target face, a new face was displayed in a different and random location. Regardless of the experimental context (e.g. background scene, no background scene), or target eccentricity (from 4 to 20 degrees of visual angle), we found that the microsaccade rate dropped to near zero levels within only 12 milliseconds after trial onset. There were almost never any microsaccades before the saccade to the face. One subject completed 118 consecutive trials without a single microsaccade. However, in about 20% of the trials, there was a single corrective microsaccade that occurred almost immediately after the preceding saccade’s offset. These corrective microsaccades were task oriented because their facial landmark targeting distributions matched those of saccades within both the upright and inverted face conditions. Our findings show that a single feedforward pass through the visual hierarchy for each stimulus is likely all that is needed to effectuate prolonged continuous visual search. In addition, we provide evidence that microsaccades can serve perceptual functions like correcting saccades or effectuating task-oriented goals during continuous visual search.

## Introduction

During active visual search, humans often make high velocity eye-movements called saccades to target various regions of interest in a scene with their gaze. When saccades are involuntary and smaller than around 1 degree of visual angle, these ballistic eye movements are typically called “microsaccades.” A recent review used scientific evidence collected over a period of more than 60 years to argue that microsaccades “are necessary to achieve continual perception during fixation” and “contribute uniquely to visual processing by creating strong transients in the visual input stream. (Rolfs, 2009).” Thus, microsaccades can play an important role in human visual perception.

One source of recent controversy was that studies investigating microsaccades generally use prolonged periods of fixation, leaving open the question of whether microsaccades occur during active visual search. That is, "microsaccades are known to occur during prolonged visual fixation, but it has been a matter of controversy whether they are also produced during free-viewing (Otero-Millan et al., 2008).” In their study, Otero-Millan and colleagues found that microsaccades not only occurred during prolonged visual search of static natural visual scenes, but also shared similar spatiotemporal properties with saccades. The authors concluded that saccades and microsaccades likely share a common neural generator (Otero-Millan et al., 2008). Additionally, their study found that microsaccades increased in frequency after a saccade landed on an area of interest in a visual scene, such as a human face.

In 1935, Buswell published a book entitled ‘How people look at pictures.’ In it, Buswell found patterns of eye fixations that were related to various patterns of elements in the pictures (Buswell, 1935; Land, 2006). In 1967, Yarbus noticed that instructions influenced these patterns of eye fixations(Yarbus, 1967). A few years later, Noton and Stark postulated a theory that humans recognize objects using small “scanpaths” in eye movements which form stereotypical patterns (Noton & Stark, 1971). This hypothesis was based on the fact that when humans look at pictures, they typically fixate in stereotypical sequences of locations that vary according to the category of the picture that was displayed. The scanpath theory was a prototypical motor theory and inspired a lot of subsequent work on perception.

How does the scanpath theory from Noton and Stark relate to the results from Otero-Milan *et al.*? In particular, are such scan-paths made with microsaccades *before* the first saccade after trial onset? If we interrupted the stimulus with a new stimulus so that a new target appeared immediately after each correct saccade entered the previous target, would the participant still attempt to explore the previous face with microsaccades? Are there any perceptual functions of microsaccades that occur after the first saccade after trial onset? If so, how long do these perceptual microsaccades take to make after the saccade landed?

To answer these questions, we analyzed a large cohort of results from a new visual search paradigm called “continuous visual search zapping” (Martin et al., 2018c, 2018a, 2018b). The “zapping” refers to the fact that as soon as the subject found the target with their gaze, that target was erased (“zapped”) and a new target was painted. Furthermore, instead of visual search on a static screen containing perhaps multiple regions of interest, this “zapping” paradigm changed the location and background scene of subsequent face targets around 18ms after the subject’s gaze reached each face. Subjects achieved up to 6.5 faces targeted each second in this paradigm (including all time for blinks and eye movements). There was not much time for fixation on any particular face because it took only an average of 18ms to paint the new target and background image.

In this paper, we explore the properties of microsaccades in this novel environment: the continuous visual search task. Based on other tasks, we formed some hypotheses of how microsaccades may behave in this task. Therefore, our work is a description of microsaccades in a new task paradigm and care should be taken when comparing our results with other papers. The purpose of the current study was to investigate whether or not microsaccades occurred during continuous visual search zapping for faces. The main hypothesis was that microsaccades would not often occur before the first saccade after trial onset. We reasoned that under a feedforward model of human visual processing, a single wave of feedforward neural activity should suffice to allow a saccade to accomplish the search task. We also hypothesized that, if we found microsaccades, that they would most often serve to perceptually correct their preceding saccades.

Nevertheless, we left open the possibility that microsaccades may occur as part of the search process, but we predicted that they would occur mainly only after saccades. Because the study from Otero-Millan *et al.* found a large number of microsaccades on areas of interest during visual search, we hypothesized that even though microsaccades would be rare, when they did occur, they would have a task-related perceptual function during successful targeting of the searched object (over-shoot/undershoot correction, face exploration). However, we hypothesized that before the first saccade after trial onset, the microsaccade rates would drop to zero like found in previous studies (Gao et al., 2015). Validating these hypotheses would be evidence that microsaccades are not critical for successful continuous visual search but do sometimes occur after saccades to more-finely hone in on the target of interest.

## Methods

### Participants

We conducted three separate experiments designed to explore the speed of continuous face detection (N1=24 subjects, N2 =24 subjects, N3=24 subjects). We did three separate experiments, but we only present the results from Experiments 1 and 3. We have numbered the experiments the same way as described in our other papers (Martin et al., 2018a, 2018c). A total of 44 subjects with normal or corrected-to-normal vision participated in a total of 72 separate sessions divided into 3 experiments: Experiment 1 (N1=24, two left-handed, 14 females, ages 21-39), Experiment 2 (N2=24, two left-handed, 10 females, ages 22-53), and Experiment 3 (N3=24, two lefthanded, 13 females, ages 21-40). Some subjects participated in more than one of the three experiments: 6 took part in all three experiments, 9 in only Experiments 1 and 2, 6 in only Experiments 1 and 3, 1 in only Experiments 2 and 3, 3 in only Experiment 1, 8 in only Experiment 2, and 11 in only Experiment 3. We recruited participants via a common laboratory mailing list for participants. Participants were compensated 15 euros for each experiment. One participant quit the task prematurely in Experiment 3, and we excluded their data and ran another participant to complete the cohort of 24 subjects in Experiment 3.

### Design

Each experiment consisted of 4 different conditions (No Scene, Upright; No Scene, Inverted; Scene, Upright; Scene, Inverted) that were separated into separate blocks. That is, in each block, participants continuously localized 500 inverted or upright faces that were either directly pasted into one of 500 different cluttered background scenes, or pasted only a gray screen as the background. Experiment 1 had a large range of eccentricities and polar angles, so that each face could appear anywhere on the screen (see Figure 1). To minimize the time required for large magnitude eye movements, every subsequent face in Experiment 3 appeared only 4° in eccentricity away from the previous face. Also, the polar angles in Experiment 3 were set such that subsequent targets appeared at polar angles of 0°-45°, 135°-235°, 315°-360° from the previous target. In Experiment 1, we pasted faces directly on the gray backgrounds or cluttered background scenes, whereas in the Experiment 3, we also locally blended the faces into the background scene by matching their grayscale histogram distribution to that of the local histogram at the pasted location (Martin et al., 2018c).

**Figure 1:**
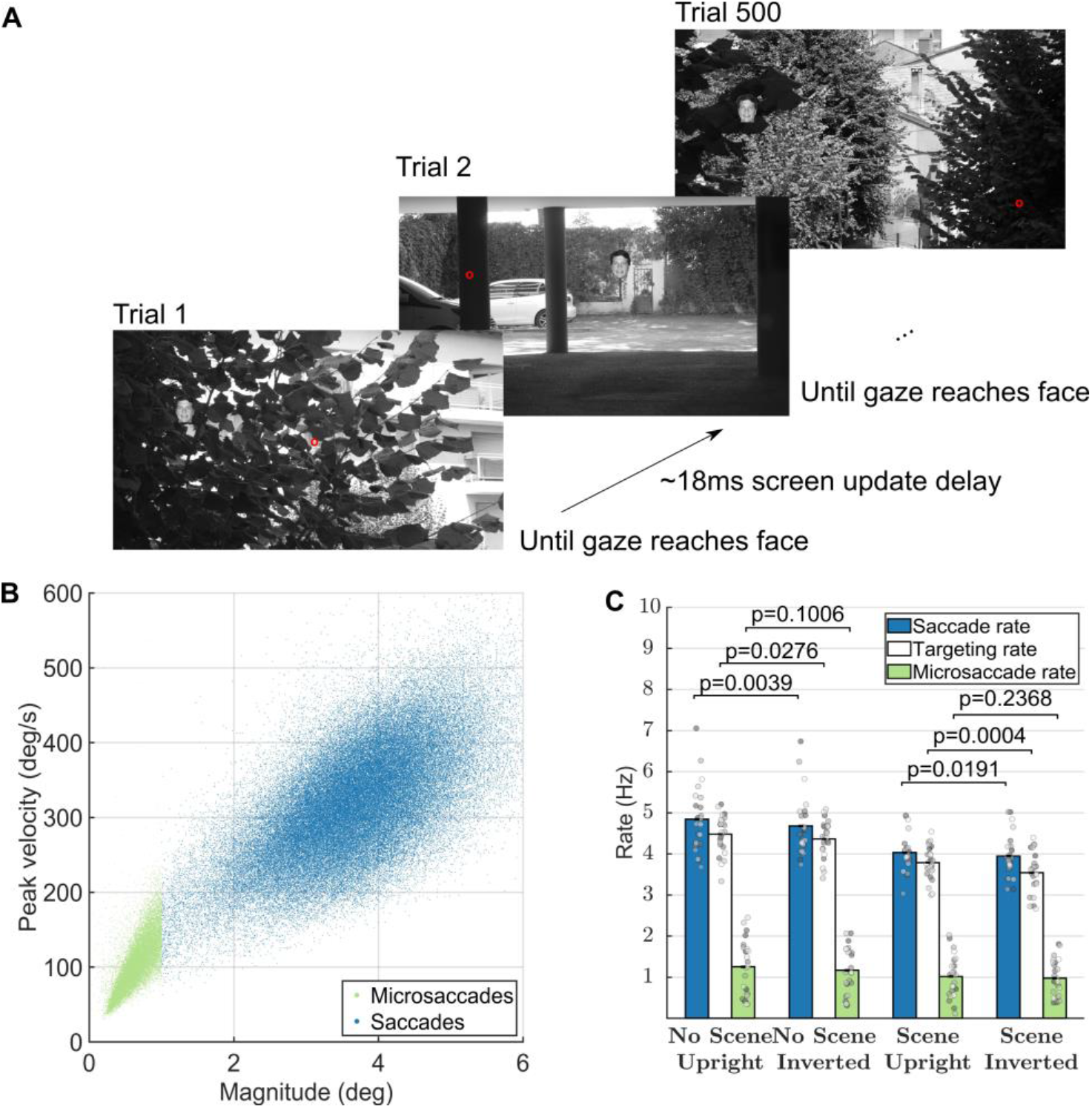
Task, presentation, and saccade analyses. (A) Task (reproduced from Martin et al. 2018). Red dot represents a hypothetical gaze location. The face shown is only for illustration, as we actually used 500 different faces in each block of 500 trials. (B) Main sequence of peak velocity versus magnitude for microsaccades (red dots) and saccades (blue dots). (C) Average subject-specific saccade, targeting, and microsaccade rates calculated within the −200ms to 800ms interval of each trial. Error bars correspond to the 95% confidence intervals around the mean. Figure 1A is an unmodified reproduction from the source’s Fig 1A (Martin et al., 2018c)

### Materials

We created the face stimuli from a set of 2316 images of segmented faces from the Humanae project (with written permission from the artist Angélica Dass, http://humanae.tumblr.com/) (Dass, 2016). Background image stimuli for all experiments were selected from a large database of 861 images, some of which have been used in previous psychophysical studies (Thorpe et al., 1996). We converted the faces and backgrounds to grayscale. We resized faces to have a height of 3° of visual angle. We resized image backgrounds to cover the entire screen resolution of 2560×1440 pixels.

Stimuli were presented on an ASUS ROG Swift PG278Q GSYNC monitor with 1ms response time and 120 Hz refresh rate, driven by two SLI linked NVIDIA GeForce 980GT GPUs, at a screen resolution of 2560×1440 pixels (Zhang et al., 2018). The display subtended approximately 31° horizontal and 22° vertical of visual angle. We controlled the display with a custom-built workstation running Gentoo Linux with a 64-bit kernel that we tuned for real-time processing. The paradigm was programmed in Matlab R2008a (The Mathworks, MA) using Psychtoolbox version 3 (Brainard, 1997; Pelli, 1997). We recorded target onset presentation times with a photodiode that was time-synchronized with the eyetracker.

We recorded eye movements using the SMI iViewX High Speed system with a 1250 Hz sampling rate. Before the first session, we determined each subject’s dominant eye and subsequently recorded and calibrated that eye. The eye-tracker sent gaze position samples at a delay of approximately 5ms to the presentation hardware. We compared the time of the entrance of the eye within the target face area and the subsequent photodiode onset for the next trial to determine that the median screen update time was 18.03ms. These values did not differ by more than 1ms when examined by condition (e.g. for Experiment 1: 17.67s, 17.95ms, 18.09ms, and 18.42ms).

To detect saccades after the experiment, we used the “microsacc plugin” for saccade detection with a smoothing level of 2 (“the raw data is smoothed over a 5-sample window to suppress noise” (R. Engbert & Mergenthaler, 2006; Ralf Engbert & Kliegl, 2003)), a velocity factor of λ=5 to determine the velocity threshold for saccade detection (“thresholds were relative to the noise level, calculated as λ = 5 multiples of a median-based SD estimator” (R. Engbert & Mergenthaler, 2006)), and a minimum saccade duration of 10 samples (corresponding to 8 milliseconds) (Ralf Engbert & Kliegl, 2003). The velocity threshold of 5 determined the velocity threshold required for saccade detection. Detection thresholds were relative to the noise level, calculated as λ = 5 multiples of a median-based Standard Deviation estimator. We considered an eye movement a saccade if and only if it went over this noise-relative threshold and had a duration of at least 10 samples, equaling 8 milliseconds (R. Engbert & Mergenthaler, 2006). Furthermore, we considered all detected saccades with amplitude less than 1 degree as microsaccades.

The parameters we chose for microsaccade detection were the same as those recommended in the detection toolbox by Engbert and used by many previous studies (R. Engbert & Mergenthaler, 2006; Ralf Engbert & Kliegl, 2003). However, note that microsaccade detection is an inexact science and recent work has seen many algorithms and toolboxes released (Nyström & Holmqvist, 2010; Otero-Millan et al., 2014). For example, decreasing the lambda parameter, which usually is set to 5 or 6, as in our study, will allow many more events to be classified as saccades (Ralf Engbert et al., 2014). To allow an exploration of the effect of different parameter choices by the community, we have described and made available the data from our study in a recent article, that researchers can freely download and explore (Martin et al., 2018b, 2018a). For example, the effects of the lambda parameter and the minimum microsaccade duration on subsequent microsaccade detection may prove a fruitful area of future research for the data in our task.

### Procedure

Each block contained 500 trials of a single condition (No Scene, Upright; No Scene, Inverted; Scene, Upright; Scene, Inverted). The orders of the blocks were counterbalanced across subjects so that each of the subjects did one of the possible 24 possible block orderings of the 4 conditions. This same order of 4 blocks was then repeated in another section of 4 blocks, so that subjects did a total of 8 blocks. After each block of 500 trials, there was a small pause of about 2 minutes while we recalibrated the eye tracker to ensure that the calibration remained accurate throughout the experiment.

Within each block, participants performed a continuous detection task in which we pasted the 3° tall face stimuli into large scenes that filled the entire 2560×1440 pixel screen of the monitor (see Figure 1A). Participants were told to find the faces with their eyes as fast as they possibly could. During the experiments, we only used the gaze position data to advance to the next trial. To make the paradigm go as fast as possible, but to also retain some robustness to noise, subjects had to maintain gaze within the 3°x3° region centered on the face for at least 2 samples (1.6 milliseconds) in order proceed to the next target. Each subsequent trial started immediately after the subject’s eye landed within the 3×3 degree window surrounding the center of the previous face (with a median screen-update error of 18ms after the subject found the previous face).

We randomly pasted the faces either based upon the position of the face in the previous trial (Experiment 3) or completely randomly within the 2560×1440 pixel scene (Experiment 1). All faces and background images within any given block were unique. We paired and combined the faces and background images before the block and counterbalanced them across subjects so that each face and background image combination appeared equally in every condition.

During the experiments, we did not place the faces based on the gaze position (which may have been better but would have slowed the experiment down considerably given the computational complexity required to blend and paste a face into a large 2560×1440 image at runtime). As subjects could have had their gaze anywhere in the 3×3 degree face detection window, the next trial’s face – which, for example, appeared at 4° eccentricity from the center of the previous face in Experiment 3 – could have been presented further or closer to the actual eye position at trial onset. Nevertheless, after the experiment, we were able to determine the actual eccentricity of the target according to the recorded gaze location at trial onset. All analyses of eccentricity used the gaze position at the time of trial onset. Even though we pasted images based on the previous target’s location, the average actual eccentricity indeed had a mean value of 4.12° during Experiment 2, and 4.09° during Experiment 3. We also checked that the average actual eccentricity did not differ by condition within each experiment (p>0.11, Wilcoxon rank sum, df=25).

## Results

Figure 1A shows the experimental paradigm, during which subjects continuously targeted small 3-degree faces. In our previous work, we did not find significant differences between the conditions in Experiments 2 and 3. Thus, the results in the current paper correspond to the data in Experiments 1 and 3 from the original datasets (Martin et al., 2018b, 2018a). Furthermore, microsaccade rates did not differ between conditions (see Figure 1C). Thus, to calculate continuous saccade and microsaccade rates and properties in Figures 2-4, we collapsed the data from all conditions. There were 24 subjects and 96,000 trials in Experiment 1 and 24 subjects and 96,000 trials in Experiment 3. Microsaccades and saccades followed the main sequence (Figure 1B). We defined microsaccades as ballistic movements with magnitude less than 1 degree (see Materials for the parameters and algorithm that we used). Saccade rates during the one second surrounding each trial were higher, but similar to targeting rates, indicating a small degree in error in targeting (Figure 1C). Microsaccade rates during the one second surrounding each trial averaged around 1Hz in each condition (Figure 1C).

**Figure 2:**
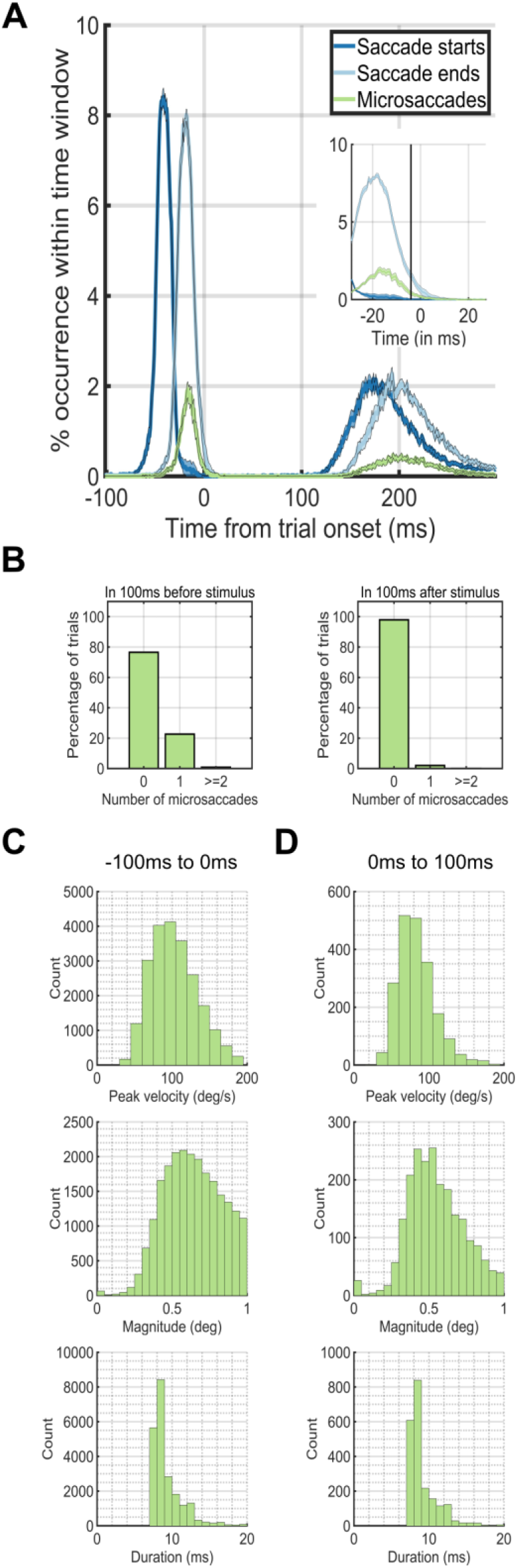
Eye dynamics around trial onset at 0ms. The peaks in occurrences of saccades before the trial onset at 0ms correspond to the preceding trial. The peaks in occurrences of saccades after 100ms after trial onset correspond to the saccades for the current trial. (A) Saccade starting times (blue) and ending times (green) plotted against microsaccade starting times (red). (B) The percentage of trials (y-axis) which had 0, 1, or greater than or equal to 2 microsaccades during the first 100ms before (left plot) or after (right plot) trial onset (0ms). (C) Microsaccade peak velocity, magnitude and duration distributions in the 100ms before trial onset. (D) Microsaccade peak velocity, magnitude and duration distributions in the 100ms after trial onset.

We next investigated the temporal dynamics of saccade rates by aligning trials based on trial onset. During the - 100ms – 0ms period before trial onset, a single microsaccade occurred in only ∼20% of the trials. On the other hand, during the first 100ms after trial onset, ∼99% of the time, there were no microsaccades at all (Figure 2A). During only 12ms after trial onset, the microsaccade rate dropped to near zero levels (Figure 2AB). We did an additional analysis that shows that the saccade-following microsaccades were not strictly always in the same direction as the preceding saccade. From the experimental data where faces were pasted using the entire screen (Experiment 1), we found that in 27% of the trials, a single microsaccade occurred directly after a saccade. Of these 27%, 84% had a polar angle target that was more than 45 degrees away from the saccadic polar angle, whereas 16% had a polar angle target that was less than 45 degrees. Thus, there were cases where the microsaccade was in the same direction of the saccade, but the subsequent microsaccade was launched more often in a different direction from the preceding saccade.

Next, we aligned the eye movement data to the onset or offset of the first saccade after trial onset (Figure 3AB). Microsaccades occurred ∼20% of the time directly after a saccade offset, but not before (Figure 3A). When we aligned trials on the saccade offset, saccades or microsaccades occurred within 25ms after the saccade offset (Figure 3B). The time before the saccade onset was almost completely without saccades or microsaccades (Figure 3C, left). Thus, the vast majority of the microsaccades we found in our study followed the offsets of the first saccade after trial onset (Figure 3D right).

**Figure 3:**
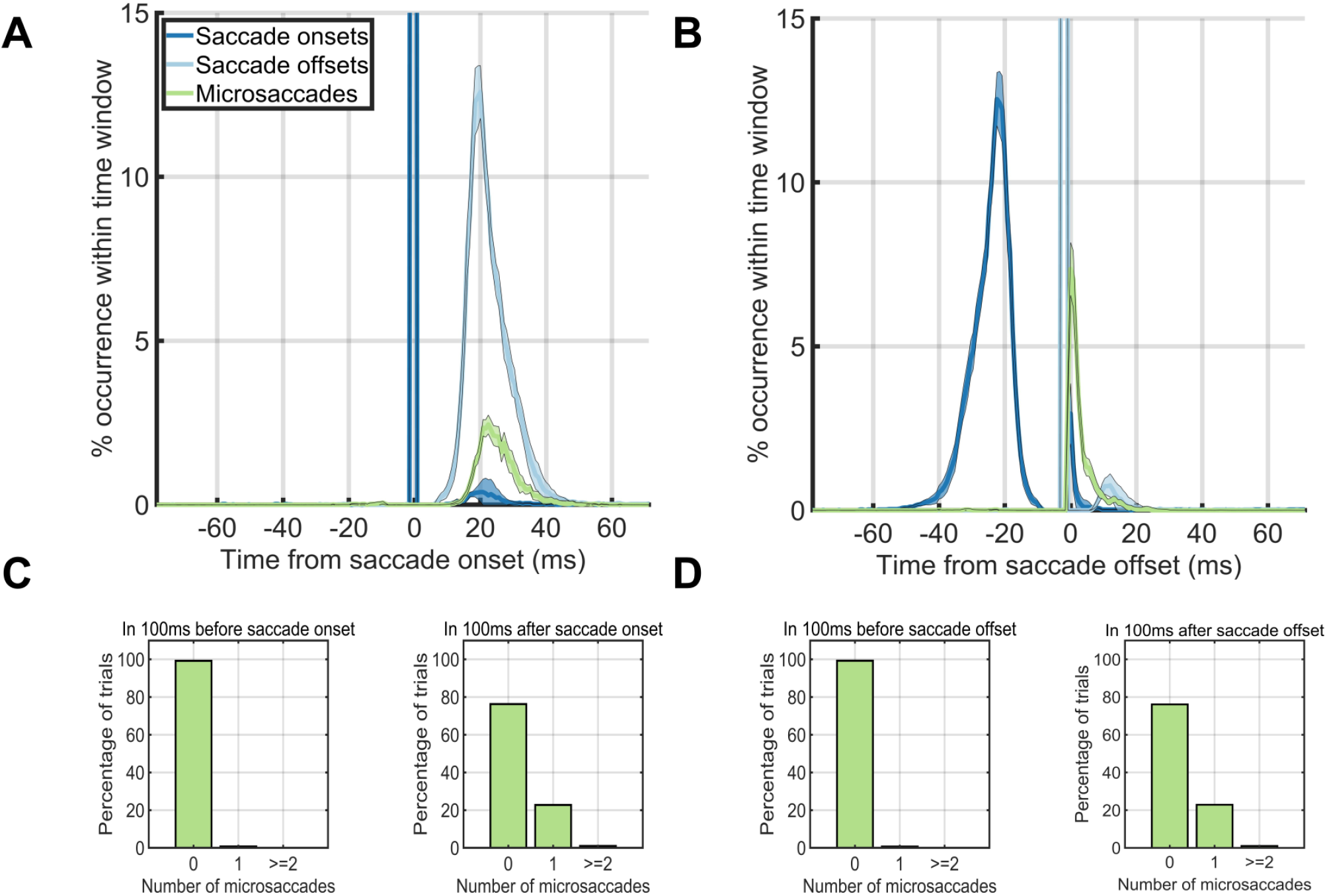
Eye dynamics around the saccade onsets (left column) and saccade offsets (right column). (A, B) Saccade starting times (blue) and ending times (green) plotted against microsaccade starting times (red) when onsets were aligned on saccade onset times (A) and when onsets were aligned on saccade offset times (B). (C, D) The percentage of trials (y-axis) which had 0, 1, or greater than or equal to 2 microsaccades during the first 100ms before saccade onset (left) or during the first 100ms after saccade onset (right) when aligned on the onset time (C) or offset time (D) of the first saccade after trial onset.

**Figure 4:**
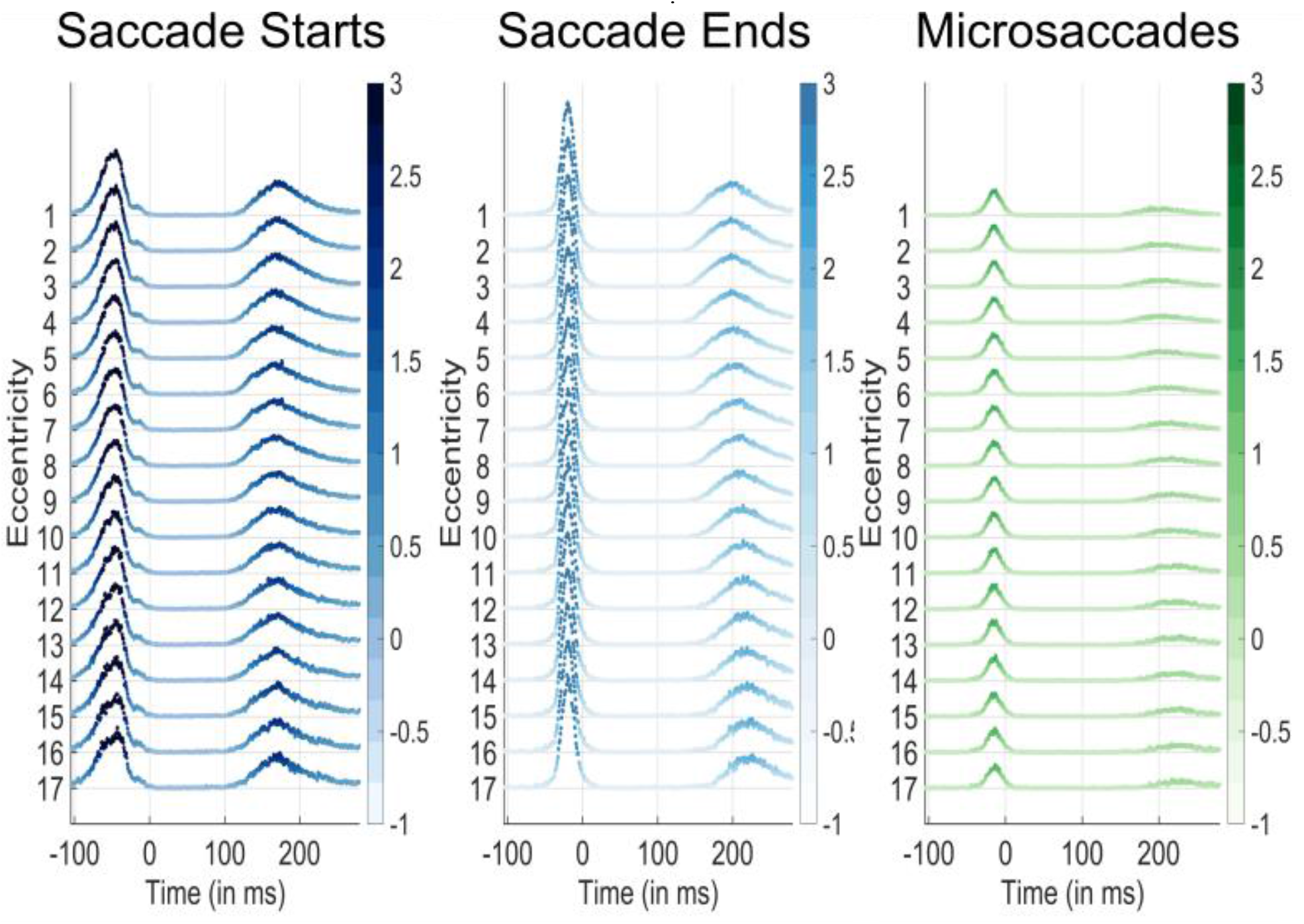
Saccade start (blue), saccade end (green), and microsaccade occurrence probabilities per time bin according to the target eccentricity of the face target that appeared at 0ms. Time 0 corresponds to the new trial onset, which was immediately after the participant’s gaze had reached the previous trial’s face. Eccentricities are listed on the y-axis. The color and height of each data point represent the percent occurrence over all 96000 trials for saccade start times (left), saccade end times (center), and microsaccade start times (right). The percent occurrence over 96000 trials is color coded according to the respective colorbars to the right of each plot. Note that microsaccades almost never occurred before the initial saccade after trial onset at any eccentricity

Figure 5A shows an example trial where there were 3 microsaccades after an initially correct saccade. It is important to note that the appearance of 3 microsaccades, like in this figure, was extremely rare. We analyzed the landing locations of each saccade and microsaccade by first recording various facial landmarks in every trial (hair top middle, forehead top middle, left eye, right eye, nose tip, mouth center, chin bottom center, Figure 5A). Next, we calculated the frequency of occurrences of saccade and microsaccade landing locations relative to the facial landmarks of a template face. We only analyzed the saccades that landed within the 3×3 degree face target area. We only counted microsaccades that started and ended within the 3×3-face target area. Concretely, for each of the aforementioned saccades and microsaccades, we linearly transformed its end point to a template face using the Procrustes algorithm (Stegmann & Gomez, 2002). Note that this algorithm does not produce an exact match to the template, because the relative facial landmarks differed from face to face. Finally, we counted the number of times a microsaccade landed at each particular location on the mapped template face and smoothed the result with a 5×5-pixel disk spatial averaging filter that subtended 0.077×0.077 degrees of visual angle. The landing locations of saccades and microsaccades had similar facial location preferences depending on whether the face was upright (Figure 5B) or inverted (Figure 5C). These results match with what has been found in previous studies of eye movements on face stimuli (Xu & Tanaka, 2013). Saccades with following microsaccades were less accurate than saccades without following microsaccades (see Figure 5CD). However, the microsaccades that occurred after saccades shared the same landing endpoint distributions as saccades without microsaccades (see Figure 5CD). Figure 5B shows that the velocity profiles were different between saccades with a following microsaccade saccade and those without a following microsaccade before the saccade offset at 0ms. We ran a temporal RUSBoost classifier at the single trial level by training on the pre-microsaccade eye velocities and trying to predict whether each saccade had a subsequent microsaccade. We found a ROC area of 0.70 (10-fold cross validation, default RUSBoost parameters, correct-trials, upright, background trials only).

**Figure 5:**
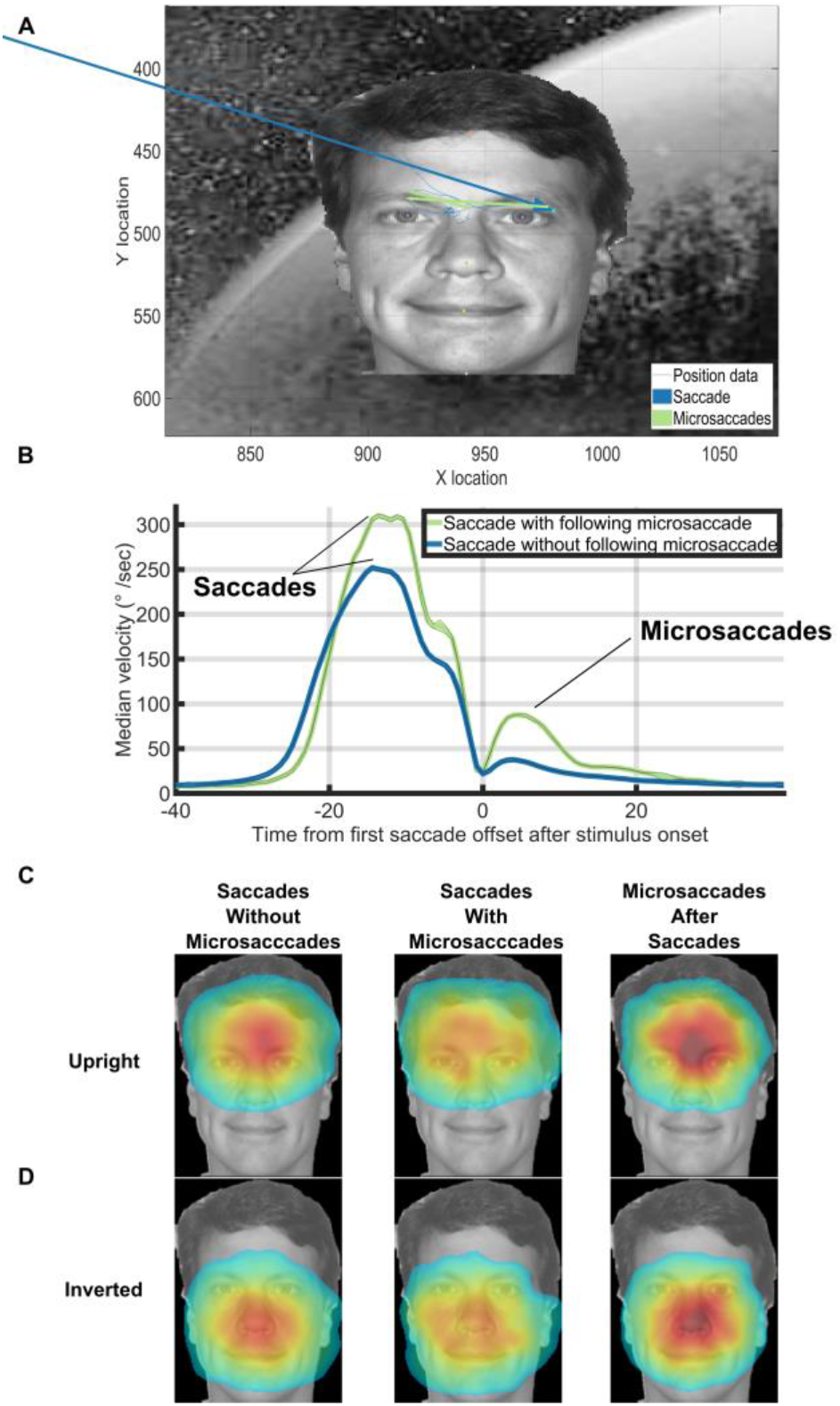
(A) An example with 2 microsaccades directly after a correct saccade in Experiment 3. (B) Median velocity on correct upright background trials at each timepoint. Trials were aligned on the offset time of the first saccade after trial onset. Note the microsaccades that occurred almost immediately after the preceding saccade (green line after 0ms). (C,D) Saccades without microsaccades and saccades with microsaccades shared the same facial landmark target frequencies within correct background trials for both upright and inverted face conditions. Saccades with following microsaccades were qualitatively less accurate than saccades without following microsaccades. However, the microsaccades that occurred after saccades shared the same landing endpoint distributions as saccades without microsaccades. The heatmaps for the inverted faces were flipped vertically for ease of presentation. To make the heatmaps, the 7 face landmark locations for each trial were mapped to the 7 face landmark locations of the template face using the Procrustes algorithm. We then mapped the first saccade after trial onset or the subsequent microsaccades into the template face’s space. Finally, we fit a normalized 2D gaussian to the raw saccade/microsaccade endpoint data using the heatmap function in PyGaze. Source images in Figures 1ACD are from one of the authors of this paper.

The results from Experiment 3 (wherein faces were only presented 4 degrees from their previous target) were similar to those in the other experiments. However, Experiment 1’s design also allowed us to confirm whether or not there was a relation between the target eccentricity and the microsaccade rate (because in that experiment, targets could appear at any location on the screen). Figure 4 shows there was no difference between the likelihood of a saccade onset or microsaccade onset across eccentricities from 4 degrees to 20 degrees. These results indicate that the microsaccade onset rate or timing did not change according to the target eccentricity.

We also calculated the number of consecutive correct trials of each possible length, ignoring sub–sequences (e.g. a run of 3 correct trials did not change the totals for runs of length 1 or 2, Figure 6A). An incorrect trial was defined as one where the first saccade after trial onset did not land on the face. A correct trial was defined as one where the first saccade after trial onset did land in the 3×3 degree square around the face. We did a similar analysis to compute the number of consecutive trials without a microsaccade (Figure 6B). The longest consecutive sequence of correct trials by any subject was 219 consecutive correct trials in the no scene, upright face condition. The longest consecutive sequence of trials without any microsaccades within the −200ms to 800ms intervals surrounding the trials by any subject was 118 consecutive trials in the scene, upright face condition.

**Figure 6:**
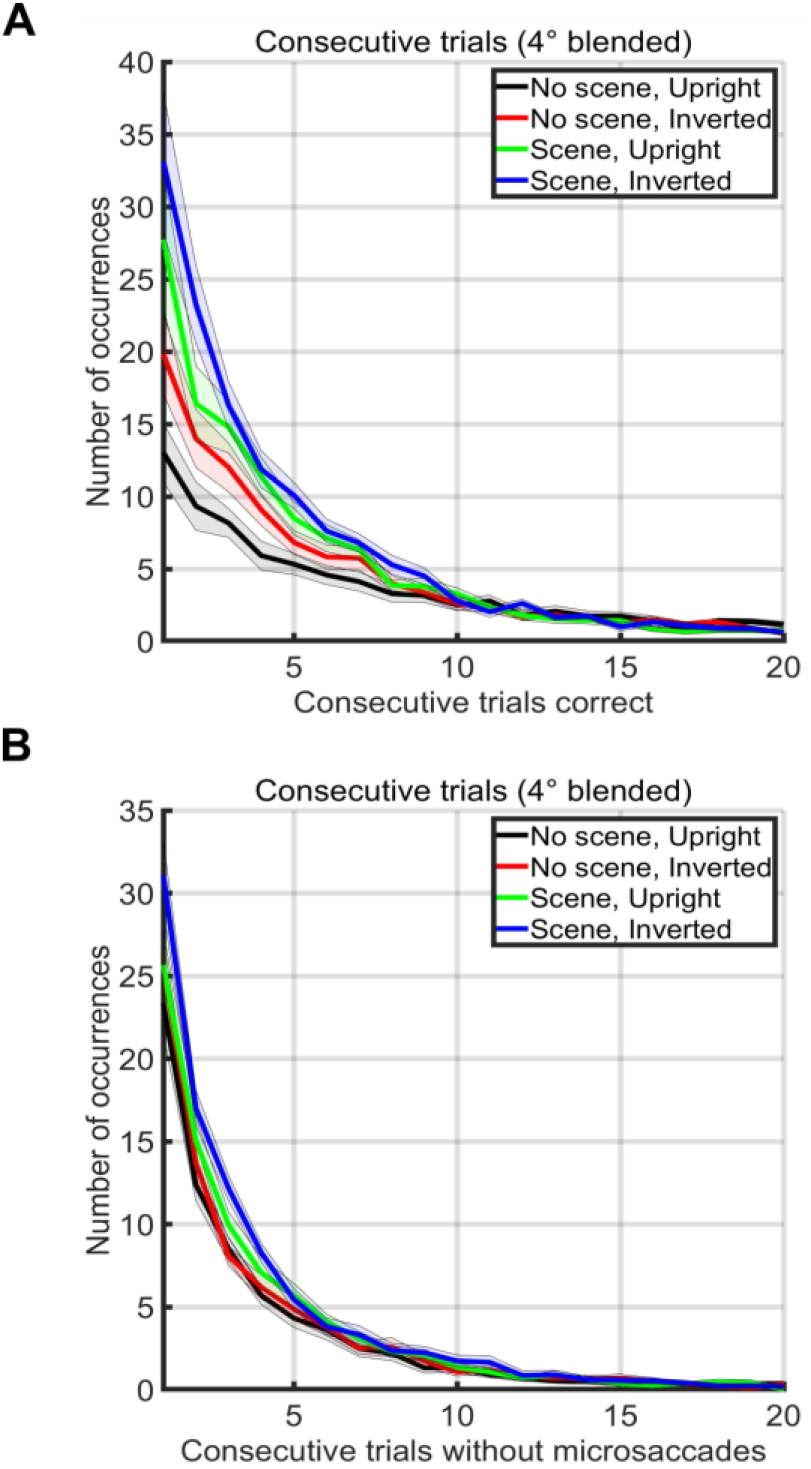
(A) Number of occurrences of consecutive sequences of correct trials without a mistake (x-axis) during two blocks of 500 trials in each condition (see legend). Note that sub-runs of consecutive trials were not counted (e.g. a run of 3 correct trials did not change the totals for runs of length 1 or 2) (B) Number of occurrences of consecutive sequences of trials without a microsaccade. Note that sub-runs of trials without microsaccades were not counted (e.g. a run of 3 trials did not change the totals for runs of length 1 or 2).

An exploration of the time required for a microsaccadic search strategy also proved illuminating. Gao *et al.* recently showed that microsaccade rates on non-continuous tasks (i.e., tasks that have pauses in between trials) drop dramatically (e.g., to less than .01 microsaccades/second) during the first 200ms after trial onset and are modulated according to task difficulty (Gao et al., 2015). Average overall rates found in such non-continuous tasks are typically between 1-2 microsaccades each second. On the other hand, we found that subjects could target up to 5-6 faces each second –rates that are 3-6 times faster than the microsaccade rates published in the literature. Therefore, it does not seem likely that the subjects could have used a microsaccadic strategy during our study and yet achieved such high rates of search. To be certain, we counted the number of microsaccades that occurred during the first 200ms after trial onset. In Experiment 3, 97.96% of trials did not contain a microsaccade in the first 200ms after trial onset, 1.98% of the trials had one microsaccade, 0.05% of the trials had two microsaccades, and 0.01% of the trials had three or more microsaccades. In addition, the average saccade amplitude after trial onset was 4.0031°. There were only 0.01% (i.e. 89/96000) of trials in which there were microsaccades away from the target during the first 200 ms after trial onset in Experiment 3. Of the subsequent saccades after these microsaccades, 84.27% landed on the target. However, of all saccades detected, 84.37% of them landed on target. Thus, subjects rarely used a microsaccadic search strategy during the paradigm, and if they did, the strategy did not yield any improvement in accuracy over a single directed saccade.

## Discussion

We were interested in finding out whether microsaccades occurred during continuous visual search. If they do exist, do the serve any perceptual function? In addition, are microsaccades critical to the success of a continuous search process? If they occur, is there any perceptual function to them or not? To answer these questions, we analyzed the properties of microsaccades that occurred during a continuous visual search task for small faces. We called this paradigm “continuous visual search zapping” to reflect the fact that as soon as the gaze reached the target, the current target was “zapped” (erased) and an entirely new scene and target was painted (Martin et al., 2018c).

Due to the high speeds of saccadic targeting during this task (Martin 2018) and the published low microsaccade rates in the literature (Gao et al., 2015), it seemed that there would not be enough time for a local microsaccadic search strategy before each saccade. Indeed, microsaccades mainly occurred only after the targeting of the face and before the next target’s trial onset (Figure 2A, 3A, 3B). During the period before saccade onset, there were almost never any microsaccades. As soon as the next target appeared, the microsaccade rate dropped to near zero levels within only 12ms. Our paradigm is different from “visual search” and “visual exploration” because the stimulus changed as soon as the eyes reach the face. We call this paradigm “continuous visual search.” It is true that subjects may have done some “visual exploration” and “visual search” until the target was found, but this was not indicated by any saccadic or microsaccadic eye movements before the first saccade after trial onset in our task. Indeed, subjects made a saccade towards the target with almost zero microsaccades intervening between the trial onset and the first saccade. Thus, there was no evidence of “visual search,” in either the saccadic or the microsaccadic sense after trial onset until the first saccade after trial onset was launched. Furthermore, microsaccades mainly occurred at the end of a saccade. Microsaccades only rarely occurred before the start of a saccade. As proof that search could successfully continue without any microsaccades at all, one participant successfully targeted up to 119 faces *consecutively* without the presence of *any* microsaccades. Therefore, it does not appear that microsaccades were a necessary component of continuous visual search and targeting.

While there were microsaccades that occurred during our task, when they did occur, they most often directly followed a saccade. These saccade-following microsaccades were task oriented. That is, such microsaccades were either corrections for an overshoot or undershoot, or local explorations within the face area. Indeed, when microsaccades occurred inside a target area after a saccade reached the target, they most often landed closer to the eyes/forehead when the face was upright and closer to the nose/chin for inverted faces (see Figure 5BC). These results are similar to those seen while recording gaze preference in previous free-viewing face inversion studies (Xu & Tanaka, 2013). Moreover, the endpoints for microsaccades and saccades both shifted their preferred locations equivalently depending on whether or not the face was inverted. Thus, the microsaccades that occurred directly after a saccade offset shared clear behavioral properties with the saccades. This means that the microsaccades and saccades both served the same perceptual function for completing the task. Interestingly, these corrective microsaccades occurred only ∼1-25ms after a saccade offset (see Figures 2A, 3AB, and 5B). This suggests that these perceptual saccade-following microsaccades were involved in some kind online mechanism that quickly corrected previous saccadic targeting attempts. These microsaccades saccades happened so fast, that they likely were planned. A very interesting question for further research is whether these microsaccades were planned: before the first saccade, during it, or immediately after checking some “hypothetical motor execution program.” Our hunch is that these microsaccades were all planned beforehand, so that the humans in our study had likely planned 12 eye movements a second, not only 6.

A previous study found that microsaccades occur during *self-paced* visual search and free viewing tasks (Otero-Millan et al., 2008). That study concluded that microsaccades have similar statistics to those of saccades and therefore likely share a common generator. This landmark study used self-paced paradigms in which the subject freely made saccades on a static screen. In contrast, our study erased the stimulus very quickly, on average only 18ms after the eye crossed into the target zone. Like the Otero-Millan *et al.* study, the microsaccades that occurred in our study mostly occurred immediately after a saccade. These microsaccades were goal oriented insofar as they landed on the same unique parts of the faces within both upright and inverted conditions as saccades (see Figure 5). Thus, the small inspection or correction microsaccades in our study were behavioral and were not distinguishable from saccades in any sense other than their magnitude or their frequency of occurrence and speed of onset. Our data therefore support the hypothesis that other studies have made that microsaccades and saccades may share a common generator (Martinez-Conde et al., 2013).

Another study concluded that microsaccades and small “inspection saccades” have fundamental differences in intersaccade intervals and generating processes (Mergenthaler & Engbert, 2010). Their study compared microsaccades in a 10-second free viewing tasks to those that occurred during a fixation task. In our study, we found almost no microsaccades during the only times in which subjects made fixations (i.e. when the previously targeted face disappeared and the next targeted face appeared). The differences between our study and the former is likely due to the continuous nature of our task, which forced the processing of an entirely new scene and target almost immediately after each saccade. Furthermore, because subjects had no way to predict the exact location of the next face target, the parallel processing “carwash” model of visual search was not likely a major factor for the saccades in our study (Wolfe, 2003, 2012a, 2012b). However, microsaccades may have indeed sometimes played a role in such a pipelined “carwash” scenario, as indicated by the task-oriented microsaccades that sometimes occurred immediately after saccade offset. Such a carwash model could have played a larger role in previous studies on microsaccades (e.g (Otero-Millan et al., 2008)), because in those studies, the stimulus was not erased when the saccade landed on the target.

Noton and Stark’s scanpath theory was a prototypical motor theory and inspired a lot of subsequent work on perception (Noton & Stark, 1971). Our data speak to this long-standing idea in the literature that the sequence of eye movements is a major part of the process of perception. Clearly, at least in our study of continuous visual search, the participants did not use local or remote scan paths to locate the targets before they launched each saccade. This fact does not preclude the possibility that scan paths may happen *after* the saccade landed on the face, but by then, the brain had already computed where the face was located.

On the other hand, after the first saccade after trial onset had already landed, about 25% of the time, there was a subsequent microsaccade (see Figure 2A). This following microsaccade was also task-oriented because it served to target the same face locations as the saccades (see Figure 5BC). In addition, in only about 20% (Experiment 1) and 27% (Experiment 2) of trials, the microsaccade was in the same direction as the preceding saccade. Thus, the post-saccadic microsaccades we found in our study were not random and did not appear to play any role for reducing retinal slip or fading. Rather, these microsaccades served a perceptual and task-oriented function. Interestingly, these perceptual microsaccades occurred very quickly after a saccade (0-25ms). Thus, they appeared to be an online redirection of saccades, perhaps indicating that the microsaccades in our study had a different generator than the saccades.

Our data fit with a whole range of “feedforward” computational neuroscience models of visual processing that use variants of convolutional neural networks (He et al., 2014; Krizhevsky et al., 2012; Riesenhuber & Poggio, 1999; Taigman et al., 2014). These experiments imply that humans should not need a lot of eye movements to recognize and localize stimuli. Indeed, once training has taken place in a deep learning system, only a single feed-forward pass is necessary to recognize just about anything (Krizhevsky et al., 2012). Consider that each saccadic movement provokes a new wave of spikes that traverse the visual hierarchy (Delorme & Thorpe, 2001; VanRullen & Thorpe, 2002). If participants can get the answer and the localization of the target on the first feed-forward pass through the visual system, then there is really no need for them to keep processing with microsaccades. That is, if subjects could compute the answer using only a single feedforward pass, then there would be no need to make a microsaccade to provoke another “wave” of feedforward activity. In effect, once they had the answer in one feedforward pass through the visual hierarchy, participants could simply make a saccade towards the target face. In our experiment, participants used microsaccades to continue to process the images when they made an error, but microsaccades did not occur during the fixation period before the saccade. Thus, the zapping task and our analyses are proof that microsaccades are not required for that first feedforward pass through the visual system. The Deep Learning and Convolutional Neural Network community, who can do just about anything on the first pass through the system, should be happy with this result.

While we did not find evidence for traditional fixation based microsaccades before the first saccade after trial onset during our task, we cannot generally invalidate the need for microsaccades during other visual tasks. Indeed, as Ditchburn (1980) wrote: “Some acrobats walk on their hands with amazing agility and most young people can learn to do this tolerably well. Certain tasks, such as following a line marked on the floor can be performed with reasonable accuracy. Yet no one suggests, from these facts, that it is mysterious that feet have evolved. Similarly, the fact that many subjects can perform certain kinds of visual tasks in the absence of frequent [micro]saccades does not conflict with the view that [micro]saccades play an important and, indeed, essential part in normal vision.” Nevertheless, it is quite impressive that the humans in our study could continuously target up to 119 consecutive faces in only around 20 seconds without the occurrence of a single microsaccade. It is important to note that our results do not invalidate any particular study of microsaccades. Microsaccades will almost certainly be very important when one needs to keep attending to a particular location (e.g. when awaiting a subtle change in some feature, for example). Our results show how humans can “walk on either their hands or feet,” with or without microsaccades, while targeting small faces in large scenes at high rates. In conclusion, microsaccades during continuous visual search did not appear to be a critical or necessary component for success in the type of continuous visual search and targeting that we used in our study. However, sometimes after a saccade, a post-saccadic microsaccade did serve a perceptual function that was task-related. Thus, microsaccades sometimes served a perceptual function after saccades, but hardly ever occurred directly before the first saccade after trial onset.

## Ethics and Conflict of Interest

The author(s) declare(s) that the contents of the article are in agreement with the ethics described in http://biblio.unibe.ch/portale/elibrary/BOP/jemr/ethics.html and that there is no conflict of interest regarding the publication of this paper. The Committee for the Evaluation of Ethics of INSERM (CEEI Institutional Review Board) approved the experimental procedures and protocol, and we obtained written informed consent from all participants prior to each experiment. The experiments were performed in accordance with all relevant guidelines and regulations.

## Acknowledgements

Funding was provided by NEI R01EY024161, ANR-13-NEUC-0004, and ERC Advanced Grant No323711 (M4). Work supported by grant ANR-13-NEUC-0004-01 (NeCiOL Project) as part of the National Science Foundation (NSF, USA), the National Institute of Health (NIH, USA) and the Agence Nationale de la Recherche (ANR, France) NSF/NIH/ANR Collaborative Research in Computational Neuroscience. NEI R01EY024161, ANR-13-NEUC-0004, and ERC Advanced Grant No323711 (M4). Copyright © 2020, the authors.

